# Distribution and genetic diversity of South Florida *Tephrosia* shed light on past cultural use

**DOI:** 10.1101/2023.01.24.524594

**Authors:** Eric JB von Wettberg, Jimi Sadle, Ezgi Ogutcen, Jennifer Possley, James Lange, Noelia Carrasquila-Garcia, Peter L. Chang

**Author notes:** Corresponding Author, 63 Carrigan Drive, University of Vermont, Burlington VT 05405.

## Abstract

The genus *Tephrosia* (Fabaceae), the hoary peas, contain high levels of rotenone, which has a long history of human use as a fish poison. We examine the distribution of *Tephrosia angustissima*, in South Florida to clarify patterns of genetic relatedness and shed light on human plant movement before European contact. Several populations of *Tephrosia angustissima* with a history of taxonomic uncertainty exist in South Florida and the neighboring Caribbean Islands.
To clarify relationships in this group, and to elucidate the conservation status of populations in Everglades National Park and Big Cypress National Preserve, we used restriction site associated DNA sequencing (RAD-SEQ) on 94 samples from South Florida and three locations in southwest Puerto Rico.
Analysis of variation in SNP markers by the Bayesian STRUCTURE algorithm and principal coordinate analysis both separated the samples into three groups. These three groups were likely separate colonization events of Florida. Genetic diversity is moderate in all of the groups, with only limited evidence of a bottleneck in some of the disjunct South Florida populations.
Overall, the human association of this group is consistent with a history of human use, suggesting conservation efforts for these taxa should consider their pre-Columbian human associations.

**Societal impact statement:** A great many endangered plant taxa exhibit patterns of edaphic specialization, occurring on particular substrates such as karst or serpentine soils. Human activities, such as the construction of shell middens, can create edaphically unique substrates. In the Americas, post-Columbian land use changes coupled with extensive loss of indigenous cultural knowledge, has created areas where associations of cultivated plants with human-generated habitats may be lost. Here we use population genetic approaches to examine rare *Tephrosia* (hoary pea) taxa from South Florida, a group of plants that produce rotenone that has been used by many indigenous groups as a fish poison. We find evidence of multiple introductions from the broader Caribbean region and an association with anthropogenic habitats such as shell middens. In efforts to conserve rare hoary peas in Florida, an understanding of past use of the landscape by native Americans is essential.

## Introduction

In an era of rapid global development, there has been a decline of indigenous land use practices. Across the Americas, indigenous land use radically altered landscapes, from the *Terra preta* “black soils” of central Amazonia to the North American great plains (e.g., Denevan 1992; Mann 2005). With the decline of these land use practices over the past few centuries, many plants that had been cultivated as food, fuel, fiber, or medicine, now face a range of pressures on their populations. Although conservation often focuses on preserving unimpacted habitats, preserving anthropogenic habitats may be essential to their persistence.

Many tropical legumes have an indigenous history of human use as fish poisons and as traditional medicines and are now declining as fish poisoning is practices less often or are outlawed. Several legumes produce rotenone, a type of isoflavanoid, that can be effective as fish poisons. When added to small water bodies, these compounds can stun fish, allowing for easy collection. In commercial applications, species of *Lonchocarpus* are most widely used, but species of *Pachyrhizus, Derris* and *Tephrosia* have similar chemical profiles. Several *Tephrosia* species from Asia, Africa, Australia as well as the tropical Americas have been noted as fish poisons, including the pantropical *Tephrosia purpurea* and African *T. vogelii* (Howe 1930, Chevalier 1937, Quigley 1956, Sauer, 1968). *T. sinapou* was a widely used fish poison by Amerindians in Guyana, although less so today now that fish poisoning is illegal and that traditional hunting practices are being lost (van Andel, 2000). This species also likely has traditional medicinal uses, and possibly was used as a soap, although these uses are not well-known (van Andel, 2000). Beyond fish poison, Speck noted that the Catawba people used *T. virginiana* to treat rheumatism (Speck 1937). Austin (2004) describes a variety of medicinal uses of *T. virginiana* by indigenous groups in the Southeastern US. (Austin, 2004).

Narrowleaf hoary pea (*Tephrosia angustissima*) is a Florida endangered species with a very narrow distribution restricted to southern Florida and Cuba. The flowers of *T. angustissima* have glabrous styles, which distinguishes this species from all other members of the genus in Florida. Three varieties of *T. angustissima* are currently considered to have occurred in Florida. Two varieties, *T*. *a*. var. *angustissima* and *T*. *a*. var. *corallicola* occurred in pine rockland habitat of Miami-Dade County, although *T*. *a*. var. *angustissima* is currently considered extinct (Gann et al. 2002). The third variety, *T*. *a*. var. *curtissii*, is known from coastal strand habitat along the east coast of Florida from Volusia County south to Miami-Dade County. In addition to southern Florida, *T. a*. var. *corallicola* has also been reported in western Cuba (Beyra Matos 1998). Based on a single Florida specimen collected in 1919, an additional glabrous-styled species was described as *T. seminole* (Shinners 1961). That specimen was collected from Godden’s Mission, an early European mission focused on converting Native Americans to Christianity that would have had an indigenous community associated with it. Godden’s Mission is believed to have been situated in eastern Hendry County; the specimen label described the plant as growing “on prairies” (Isely 1981, 1982). The herbarium labels for the Godden’s mission *Tephrosia* said it was used to treat nosebleeds and other maladies. A subsequent taxonomic treatment of *Tephrosia* included *T*. *seminole* as a synonym for *T*. *a*. var. *curtissii* (Isely 1981).

All three varieties of *T. angustissima* are extremely rare. In the wild, *T*. *a*. var. *curtissii* is the most widely distributed variety with several extant populations known along the eastern coast of South Florida and with a recent population estimate of 2000 plants (Wendelberger 2010, Wendelberger and Maschinski 2016). A single natural population of *T. a*. var. *corallicola* is currently known in Miami-Dade County. In addition, two populations of this variety were introduced on conservation land in close proximity to the natural population by staff at Fairchild Tropical Botanic Garden with their partners in the Miami-Dade County Environmentally Endangered Lands (EEL) Program (Wendelberger 2010, Wendelberger and Maschinski 2016, Possley et al. 2022).

The taxonomic uncertainty surrounding *Tephrosia* also limits our understanding of its historical population size and its human use. Although once more abundant, perhaps due to indigenous human use, the species may have experienced a reduction in population size in the past. It is likely that *Tephrosia angustissima sensu lato* colonized Florida from geologically older and higher land in the Caribbean. Known as a prized fish poison to Native Americans, populations may have been transported to Florida from the Caribbean instead of or in addition to natural dispersal. Irrespective of how they were introduced, populations in Florida likely underwent a severe population bottleneck upon arrival that led to reductions in population size and low genetic diversity. The loss of genetic diversity would limit the capacity of the Florida populations to respond to subsequent environmental changes, such as sea level rise and the introduction of invasive species.

Here we have developed genetic markers to resolve the relationships of different taxonomic groups of *Tephrosia* to one another and provide insight into whether the distribution of groups suggests possible past human movement of this group. Resolving taxonomic groups also allows us to define management units that are genetically similar. Furthermore, we aimed to elucidate historical demographic and migration patterns to better understand the likely consequences of ongoing population loss and the effects of potential management actions.

## Methods

### Biological materials

We first developed a list of plants to collect from a variety of distinct locations (Figure 1). Our collection sites included all populations of putative *Tephrosia angustissima* in South Florida, as well as *T. florida*, which occurs more widely in South Florida. We collected from Russell Key in Everglades National Park (a site linked to the Calusa people) and three sites within the Big Cypress National Preserve, including a location close to the likely site of Godden’s mission visited by Europeans and Seminoles. In Miami Dade County, we sampled a small population of *T. a*. var. *curtissii* from Haulover Park and a population of *T. a*. var. *corallicola* from Chapman Field, a USDA Agricultural Research Service Station. Accessions from Chapman Field are also held as an *ex situ* collection at Fairchild Tropical Botanic Garden and were used for introductions at two nearby Miami-Dade County preserves: Ludlam Pineland and the Deering Estate. At Ludlam Pineland we collected wild *T. florida* to serve as an outgroup for the clustering analysis. We were also able to obtain roadside samples of *T. cinerea* from a collecting trip to southwestern Puerto Rico in 2017. Material was obtained from three localities in Municipalidad Cabo Rojo: two more montane sites [**Sierra Bermeja:** Cabo Rojo NWR “Cerro Mariquita”, 26 Jan 2016, *Lange 16* (MAPR); ‘Finca Escabi’, 27 Jan 2016, *Lange 17* (MAPR); **El Conuco:** ‘Upper Rancho Hugo’, 28 Jan 2016, *Lange 19 w/Possley* (FTG); El Conuco ‘Finca Solins’, 28 Jan 2016, *Lange 20* (FTG)] that are fairly close together and similar and a third along road 301 near the **Cabo Rojo** lighthouse [28 Jan 2016, *Lange 18* (FTG)].).

**Figure 1.**
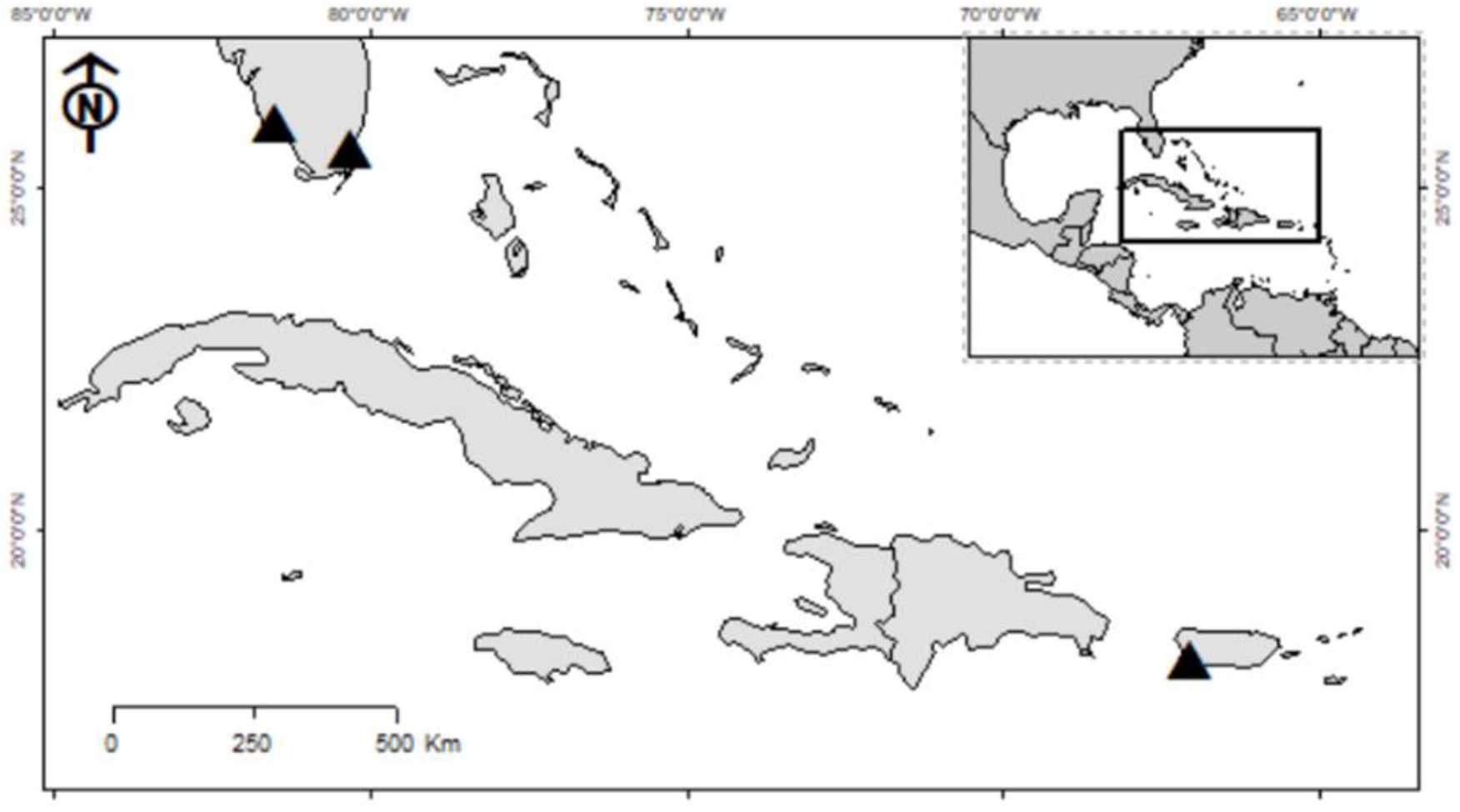
Map of the Caribbean showing sampling locations in Florida and Puerto Rico.

For sequencing, we used a slight variation on the RADseq technique, double digest RAD (ddRAD), which achieves more reproducible results with as little as 100 ng of genomic DNA (Peterson et al. 2012). DNA from leaf tissues was extracted from all collected specimens using a Qiagen DNeasy kit (Germantown, MD, USA) and extraction yields were assessed fluorometrically. The 94 samples with the highest quality DNA, representing at least 8 samples from all nine *Tephrosia* locations, were processed for restriction-site associated DNA sequencing. High quality genomic DNA (300 ng) was digested with two restriction enzymes simultaneously. Adapter sequences, P1 (containing a unique barcode and PCR primer site) and P2 (containing a PCR primer site) were ligated to the overhanging ends created by the restriction enzymes. Fragments of the appropriate size for sequencing (~300 bp) were selected using a Pippin Prep kit (Sage Science), followed by PCR amplification, during which index sequences were added. All samples were sequenced on an Illumina HiSeq 2500 Sequencing System at the UC Davis Genome Center. ddRAD sequencing reads underwent quality assessment and SNPs, genotypes, and haplotypes were called using the GATK software pipeline with recommended settings (McKenna et. al, 2010). We retained 6278 SNPS after filtering for minimal coverage (10 reads) and to remove singletons. Sequencing multiple samples from all nine locations gave us sufficient replication to compare levels of genetic variation in each location, and to infer historic population sizes. We decided against further sampling so as to not reduce sample sizes below a threshold where variation cannot be estimated.

We used Genalex 6.05 (Peakall and Smouse 2006, 2012) to calculate a range of population genetic statistics to understand patterns of variability and make demographic inferences. Estimates of expected and observed heterozygosity as well as deviations from Hardy-Weinberg equilibrium genotype frequencies were calculated to establish hypotheses concerning relationships between population size, genetic structure, and clonal propagation (e.g., Gravuer et al., 2005). The Haulover population was dropped from these analyses for a small population size. Analysis of molecular variance (AMOVA, Excoffier et al 1992) was performed to partition molecular variation among species, varieties, including variation within and between populations. STRUCTURE, a clustering algorithm, was used to assign individuals to populations and to identify admixture between populations (Pritchard et al 2000, Evanno et al 2005). A principal coordinate analysis was used as an alternative clustering approach, which was implemented in Genalex 6.05 (Peakall and Smouse 20s06, 2012). These two analyses are critical to defining management units, and circumscribing areas and genotypes that are compatible for reintroductions or augmentations without risk of hybridization above which occurred in the past.

A Treemix analysis (Pickrell and Pritchard, 2012) was performed to detect introgression among different groups. The relationship between allelic richness and gene diversity, and patterns of linkage among loci were examined for signs of bottlenecks and multiple colonizations in Florida.

We aimed to estimate current and past population sizes in two complementary ways. For effective population size, we used a program called NeCalculator2 (Do et al., 2014) to estimate effective population sizes based on the molecular co-ancestry method, since the linkage disequilibrium and heterozygote excess methods estimated infinite population sizes for these taxa. We estimated population size over the past thousands of years based on the site frequency spectrum, SFS (e.g., Ragsdale et al., 2018). RAD-seq. is an imperfect tool for calculating the absolute value of population size since the calculation depends on the number of invariant sites. This has always been an issue since that number is not really known with RAD-seq analyses (e.g., Gattepaille et al., 2013). In our processing steps, we identified sites that are segregating, but otherwise, we do not know if sites are not sequenced or if they are invariant). However, we can use the snp loci as “sites of interest”, providing us with 6278 segregating snps among 94 samples.

## Results

### Clustering algorithms

We developed 6278 SNP markers with RAD-seq, after filtering. Both STRUCTURE and PcoA consistently identified three clusters in the material, consistent with three taxa (Figure 2). At a K of 3, STRUCTURE separates *T. florida* from Ludlam Pineland into one group, the Big Cypress location into a second cluster, and Russell Key, Chapman Field, and the Puerto Rican samples into a third cluster. As interpretation of population assignment can have value at nearby Ks, we also looked at K=2 and K=4. At K=2, Big Cypress is separated from Russell Key, Chapman Field, and Puerto Rico, with the Ludlam location admixed. At K=4 the same groupings occur, but the Florida Russell Key and Chapman field populations are separated from the Puerto Rican samples. Principal coordinate analysis (Figure 3) gave a similar result. A Treemix analysis (Figure 4) showed no sign of admixture among the three taxa.

**Figure 2.**
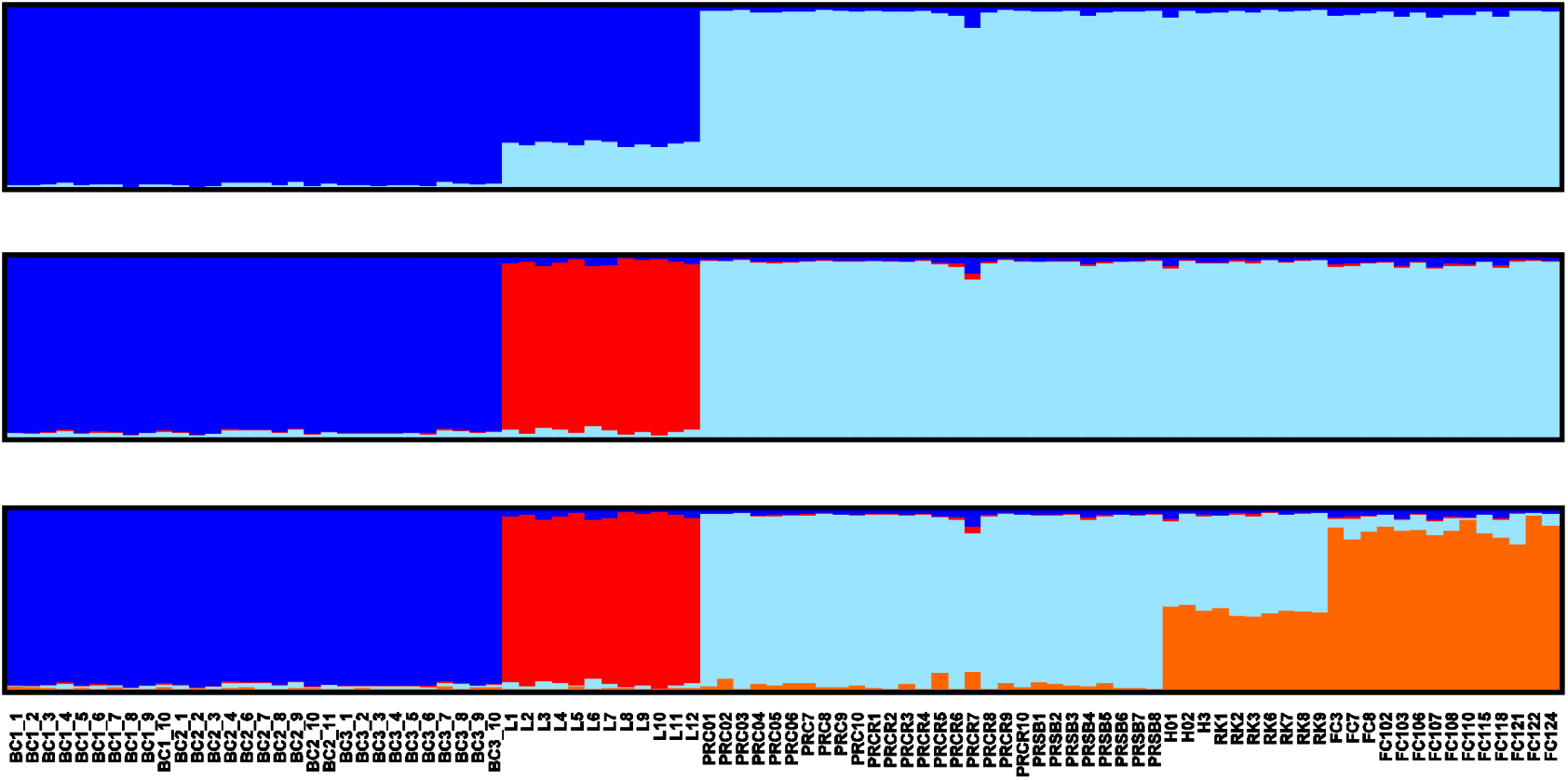
Results of a STRUCTURE analysis on 6272 SNPs, with population numbers (K) of K =2, K=3 and K=4 shown. The Evanno method determined K=3 to best describe the data. K=2 and K=4 are shown to contextualize our interpretation of assignment of samples of populations.

**Figure 3.**
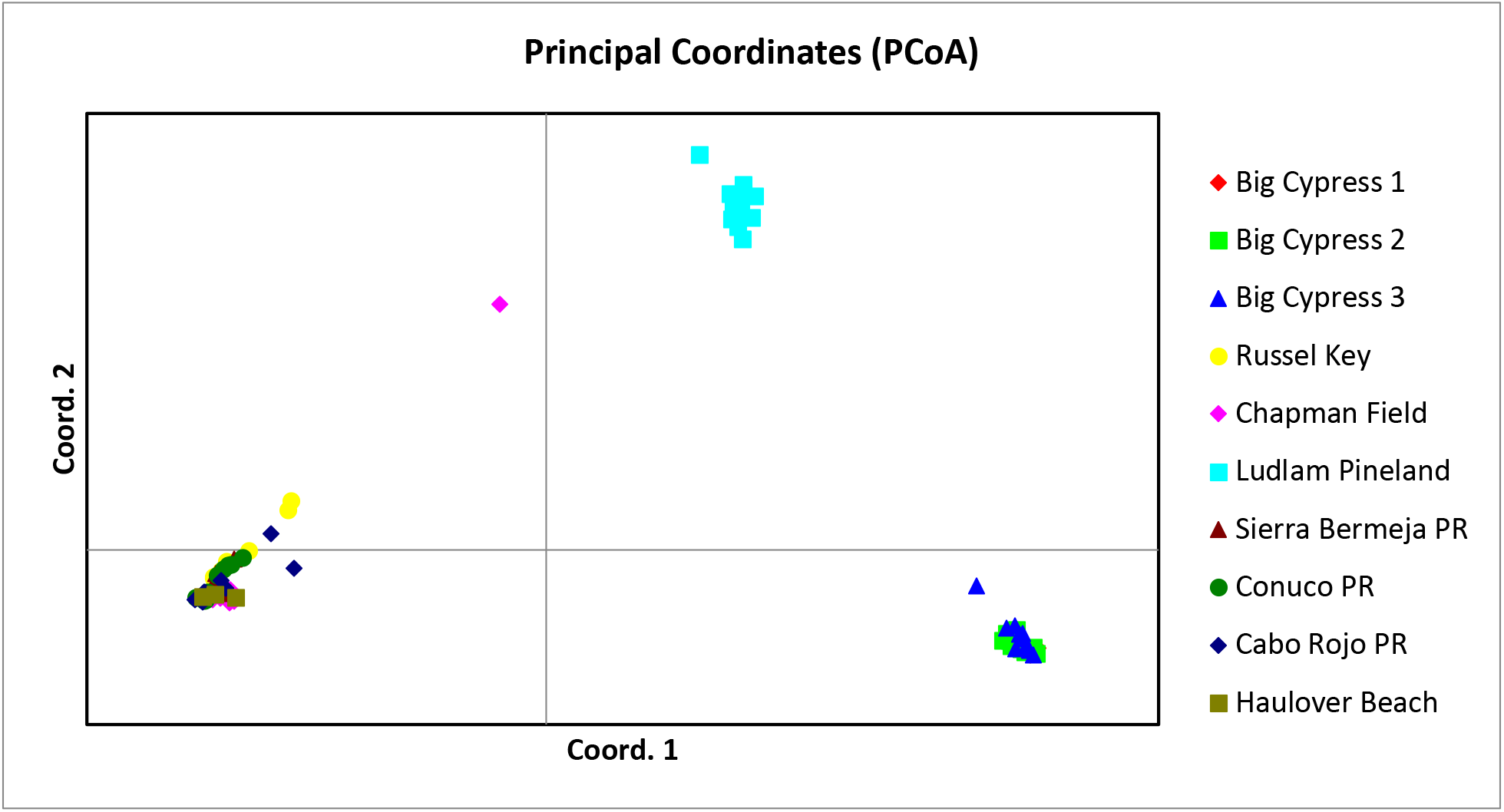
Principal coordinate analysis. Populations 1-3 are from Big Cypress (lower right corner). Population 4 is Russell Key, Population 5 is Chapman Field. Population 6 is Ludlam Pineland. Population 7-9 are from Puerto Rico. PCoA coordinate axes one and two represent 42.6 and 19.3 percent of diversity, respectively.

**Figure 4.**
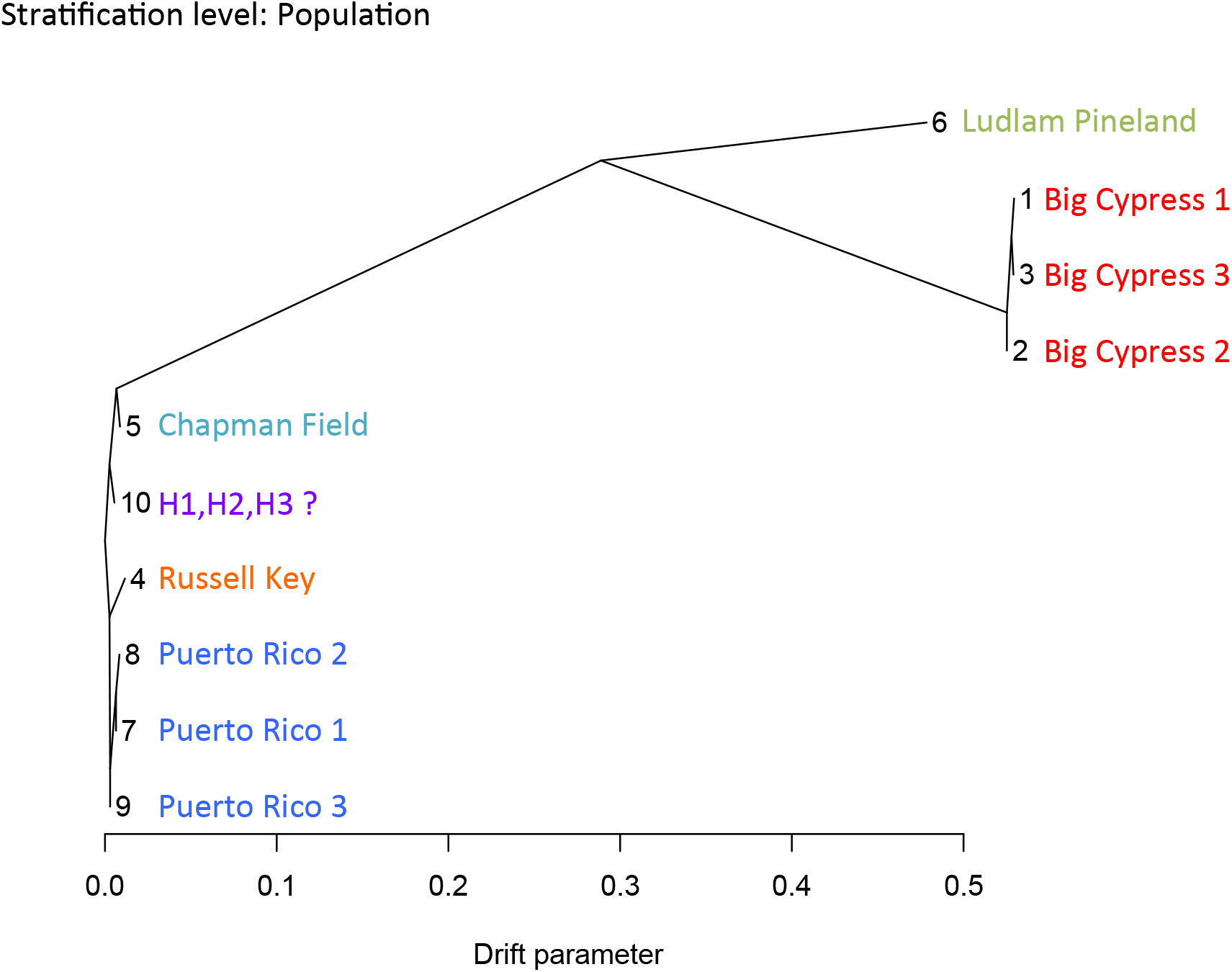
Treemix analysis showing a lack of introgression among the three South Florida *Tephrosia* taxa.

### Diversity metrics and differentiation among populations within taxa

Estimated standing diversity is similar across the 9 sampled locations, both as gene diversity (He) and as Shannon’s diversity index (Table 1). Percentage of polymorphic loci are similar across sampled locations, hovering around 80%, with the highest polymorphism rate in the one of the restored populations at Chapman Field.

Fixation index (F_ST_) was high among taxa (above 0.3) and low within taxa (< 0.1), indicating recent divergence among taxa with low diversity within taxonomic groups. However, F_ST_ was slightly higher between the two Florida *T. angustissima* locations and the three Puerto Rican locations (~0.01-0.03, Supplemental Table 1). AMOVA partitioned 21% of variation among taxa (Supplemental Figure 1), and 78% within individuals, with none among populations. This is consistent with the low within taxa F_ST_ estimates. PcoA within taxa showed similar patterns, with little differentiation among the three Big Cypress populations. Nonetheless, PcoA did detect some differentiation among the two Florida *T. angustissima* and 3 Puerto Rican *T. cinerea* populations, even when F_ST_ values were quite low.

### Effective population size and historical population size

Estimates of effective population size from NeEstimator are given in Table 2. These estimates are low with confidence intervals that overlap or nearly overlap with zero. These estimates are generally low, consistent with population size declines over the past two centuries.

The Big Cypress population (Figure 5, Blue color, T1), shows a drop in Ne (effective population size) at about 7K years ago, steadying off at Ne = 800K (Figure 5). The Ludlam Pineland *T. florida* population (T3, red color) started decreasing about 12K years ago and has been decreasing since then, only steadying off about 200 years ago. The Russel Key/Chapman Field/Puerto Rican group shows higher and consistent population size (Black color, T2). Although the Ne values should not be interpreted as numerically precise, qualitatively they are consistent with population bottlenecks around the time arriving in Southern Florida since the last glacial maxima. The Puerto Rican samples (T2) did not converge and showed little shift in population size over the past several hundred thousand years.

**Figure 5.**
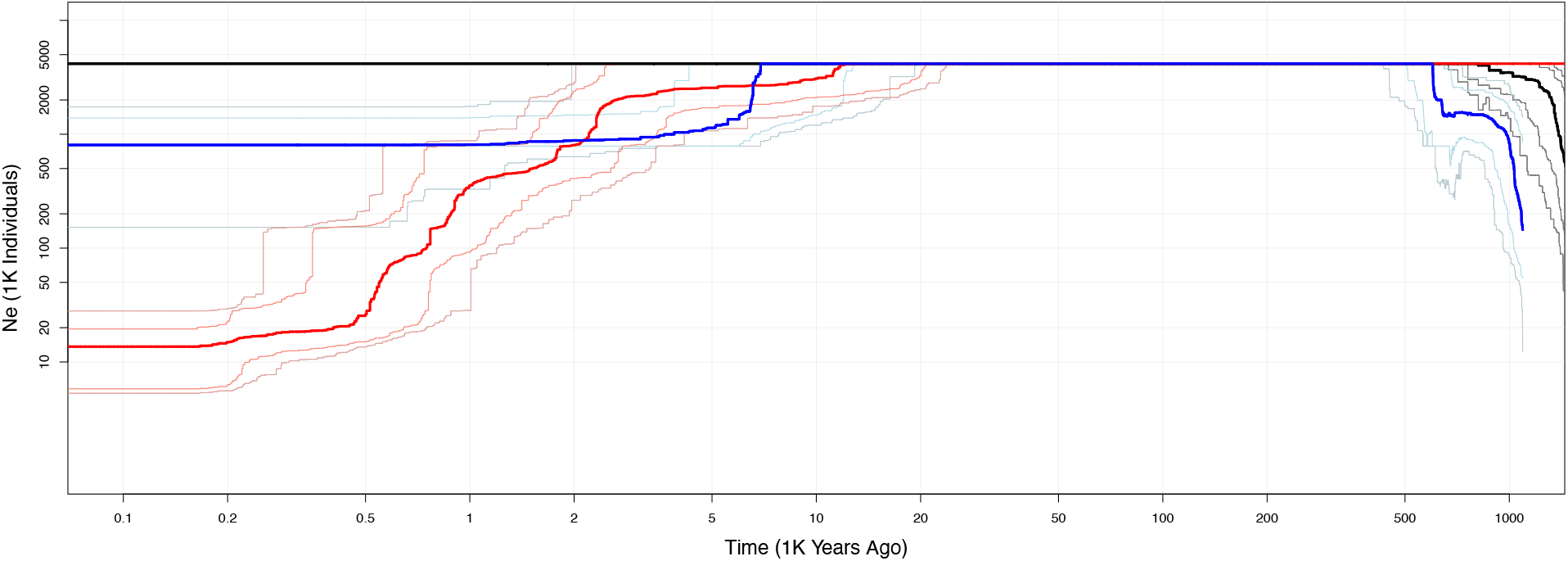
Historical effective population size estimates based on the site frequency spectrum. Blue color indicates the Big Cypress population (T1), Red the *T. florida* population from Ludlum (T3), and black the Russel Key/Chapman Field, Puerto Rican group ((T2). Solid lines represent the estimated population size, and dotted lines the 75 and 95% confidence intervals around the estimate.

## Discussion

Our sampling provides some insight into the relationship of *Tephrosia angustissima sensu lato* in South Florida. Our analyses clearly indicate that plants from Big Cypress are quite distinct from populations in Russell Key and Chapman Field as well as those from Puerto Rico. The Russell Key site is a location associated with the Calusa people, one of the pre-Columbian groups of Florida. The Big Cypress sites are close to the historical location of Godden’s mission, where trade and cultural exchange occurred between Europeans and the Seminole, a group that fled to South Florida after wars and persecution in the mid-19^th^ century. Consequently, we recognize that the current taxonomic classification of *T. angustissima sensu lato* represents at least 2 distinct taxonomic units. Due to DNA degradation issues, we were unable to get similar types of sequence data from herbarium samples. Therefore, with the apparent extinction of *T. angustissima* var. *angustissima*, we cannot directly address the subspecies question with complete confidence. However, we note that some of the differences could be due to environmental as well as genetic differences. The relatively low amount of differentiation between the Russell Key/Chapman field populations and the three Puerto Rican populations, relative to the more highly diverged Big Cypress population, are consistent with *Tephrosia seminole* being a distinct entity, and *Tephrosia angustissima* being part of a more widespread Caribbean species. These patterns are also consistent with multiple introductions of *Tephrosia* to South Florida, perhaps facilitated by separate groups of humans, although we have limited capacity to make inferences about that.

Without broader sampling of the Caribbean populations, in particular Cuba, determining the number of introductions of these taxa to Florida is not possible. Broader sampling efforts with voucher specimens are critical to unravel a more thorough demographical history of this group. We hope in the future such sampling is possible. We know that many taxa in South Florida have close relatives in Cuba, yet, at the very least our analysis suggest that the Russell Key location is genetically very similar to locations from southeast Florida Pine Rockland and coastal strand, which was unexpected. Our findings point to a potential single introduction of this taxa into Florida, that would have involved long distance dispersal within southern Florida. The presence of these taxa in both pine rocklands and coastal strands of the eastern coast of Florida and a shell mound in Southwestern Florida suggest a potential cultural link between the two regions.

Lastly, our results revealed very small effective population sizes for all *Tephrosia* locations sampled. In our collection sites, all of the sampled locations had fewer than 50 stems, indicative of very small populations that are widely isolated from one another. As a result, it is likely that these groups exhibit persistent small population sizes that are exacerbated by genetic bottlenecks during introduction to new locations. We argue that continued conservation of all these locations is essential especially for populations in unusual and threatened habitats, such as shell middens, which are of particular interest due to their disjunct distribution, human history, and unique adaptations to challenging environmental conditions. The Chapman Field site, which has been used as a seed source for reintroduction plantings, has high observed levels of diversity, and should be managed as an important germplasm repository for further reintroductions.

Native American populations in the study region reflect an unfortunate and sad history since European arrival, with near total annihilation of indigenous populations in Cuba, declines and significant cultural shifts in the Tequesta, Calusa and other groups that have been in South Florida since Spanish colonial times, and Miccosukee and Seminole groups that fled to the Everglades region following eviction and genocide during the Trail of Tears and Seminole wars of the mid-19^th^ century (Wasserman, 2009). Written records from colonial times of the Tequesta and Calusa peoples that inhabited South Florida before European contact are limited, with most of the survivors of the groups perishing when evacuated to Cuba when Florida was transferred to British rule in 1763. Consequently, ethnographic approaches that document use of *Tephrosia* as a fish poison in Florida may not be able to uncover past use. However, the distribution of these taxa is highly suggestive of a pattern of human dispersal to anthropogenic sites such as shell middens and former mission sites. Furthermore, as a taxon closely associated with indigenous human use, they likely declined as disease, 19^th^ century wars, and 20^th^ century development altered the indigenous anthropogenic habitat where they thrived.

The distribution of *T. angustissima*, which is geographically restricted in the US and includes anthropogenic sites such as shell middens and former mission sites, is highly suggestive of movement by humans. However, written records of plant use by South Florida’s early indigenous groups from colonial times are limited to a few accounts and only include a few species. Consequently, ethnographic approaches that document use of *Tephrosia*, whether as a fish poison or other use in Florida, were generally not available as a means of understanding the current distribution of this species. The approach taken in this study offers a novel means of assessing the potential role of humans in the distribution of plant species where this information is lacking. Our work demonstrated that plants collected in Big Cypress were genetically distinct when compared to the rest of the populations. Otherwise, there was not clear genetic evidence to suggest human movement in the other taxa.

## Acknowledgements

We thank the USDA Chapman Field Station, Miami-Dade County Parks, and Miami-Dade County Environmentally Endangered Lands for permission to collect samples. Yadira Reynaldo processed samples for DNA extraction. This research was supported by a cooperative agreement between Everglades National Park and Florida International University.

## Author Contribution

The project was conceived by JS and EvW, with input from JP. Funding was obtained by EvW and JS. Collections were performed by JS, EvW, JP, and JL. Laboratory work was performed by NCG, and analyses were performed by PC, EO, and EvW. EvW pulled together the manuscript, with help from all authors who wrote subsections and revised multiple versions.

## Data Availability Statement

All sequence data will be available on NCBI upon acceptance of the manuscript. Processed files are available on the open science foundation page for this project, https://osf.io/q5wzu/.

